# Monarch butterfly and milkweed declines substantially predate the use of genetically modified crops

**DOI:** 10.1101/378299

**Authors:** JH Boyle, HJ Dalgleish, JR Puzey

## Abstract

Monarch butterfly *(Danaus plexippus)* decline over the past 25 years has received considerable public and scientific attention, in large part because its decline, and that of its milkweed *(Asclepias* spp.) host plant, have been linked to genetically modified (GM) crops and the associated herbicide use. Therefore, the monarch has emerged as a poster child for the anti-GM movement. Here we use museum and herbaria specimens to extend our knowledge of the dynamics of both monarchs and milkweeds in the United States to more than a century, from 1900-2016. We show that monarch population trends closely follow those of their milkweed hosts; that both monarchs and milkweeds increased during the early 20th century, and that recent declines are actually part of a much longer term decline in both monarchs and milkweed beginning around 1950. Herbicide resistant crops, therefore, are clearly not the only culprit, and likely not even the primary culprit, as these declines began decades before GM crops were introduced.

---

Genetically modified (GM) crops have had a profound effect on agriculture, but their impact on the natural world is controversial^1^. The monarch butterfly is one of the few described instances of GM crops causing declines in a species outside of agricultural fields^1^. Monarchs’ dramatic recent decline is linked to GM crops in the scientific^2,3,4,5^ (but see^6^) and public^7, 8^ literature. Accordingly, monarchs are an emblem of the anti-GM movement, including being used in the logo of the food-labeling “Non-GMO Project” (nongmoproject.org).

The monarch butterfly *(Danaus plexippus)* is a large, showy Nymphalid butterfly best known for its migration, in which monarchs from a small overwintering area in Mexico recolonize breeding grounds across eastern North America over the course of several summer generations, followed by a single migration back to Mexico in the autumn^9, 10^. Over the past 25 years, this migratory population of the monarch has experienced a drastic decline, as much as 80%, as measured at the overwintering area in Mexico^4, 11^. Surveys of both immature and adult stages suggest a decline at the breeding grounds as well^12^. Previous work has identified the decline of milkweed *(Asclepias* spp.), monarch’s food source and egg nursery, as a likely culprit in monarch decline^3^. GM crops, in turn, have been identified as the major cause of milkweed decline^3, 13^: Because GM crops are frequently engineered to be resistant to glyphosate or other herbicides, herbicides are sprayed indiscriminately across crop fields killing all non-GM plants. This is especially harmful to common milkweed, *A. syriaca.* Although *D. plexippus* caterpillars are able to feed on at least thirty species of milkweed^14^, currently the most important host species for *D. plexippus* in their summer breeding grounds is *A. syriaca^15^*, likely because of its former abundance in agricultural fields^16, 17^.

It is clear that herbicide treatments kill milkweed; however, the importance of GM crops in milkweed and monarch declines is not yet clear, with some evidence pointing to other factors as more important drivers of the observed decline. The best evidence for this is that the decline of monarch butterflies appears to predate the use of GM crops. The monarch population size has been recorded in the overwintering grounds since 1993^11^, and the population decline is thought to be either linear or exponential over this period^2, 12^. However, herbicide-resistant crops were not introduced until 1996, and initially accounted for only 2% of US cropland (Figure S4). Herbicide-resistant GM varieties are available for corn, soy, and cotton; half of the acreage of these crops was herbicide resistant in 2004, and half of all crops were herbicide resistant by 2013 (Figure S4). Since few acres were planted with herbicide resistant crops during the beginning of the monarch decline, monarch and milkweed declines may have begun some time before the advent of herbicide tolerant crops. However, because monarchs, like many insects, exhibit substantial year-to-year variation in population size^11^, it is challenging to test this hypothesis using the currently-available data sets, which include only 10 or so data points from before the widespread use of GM crops. Here we use natural history collections to test this hypothesis across a much longer period spanning the 117 year period from 1900-2016.

## Results

### *Occurrence trends in the genus* Asclepias *and species* Danaus plexippus *from 1900-2016*

We extracted digitized collection information for over 5,000,000 insect and plant records and counted the number of specimens of *D. plexippus* and *Asclepias* collected in each year from 1900-2016 (sample sizes in Table 1). Since collection effort has varied over this time period (Figure S1), we accounted for collection effort by calculating “relative occurrence” for both groups. To do so, we divided the number of milkweed and monarch specimens collected each year by the total number of vascular plant and lepidoptera specimens, respectively, collected within the same geographic range and year. Our data set does not include records from states west of the continental divide, as these states are home to a population of monarchs that is geographically distinct from the eastern migratory population^18^. We present these data in Figure 1A, alongside a smoothed mean and 95% confidence interval, done using Loess smoothing with the default smoothing span as implemented in ggplot2^19^.

**Figure 1:**
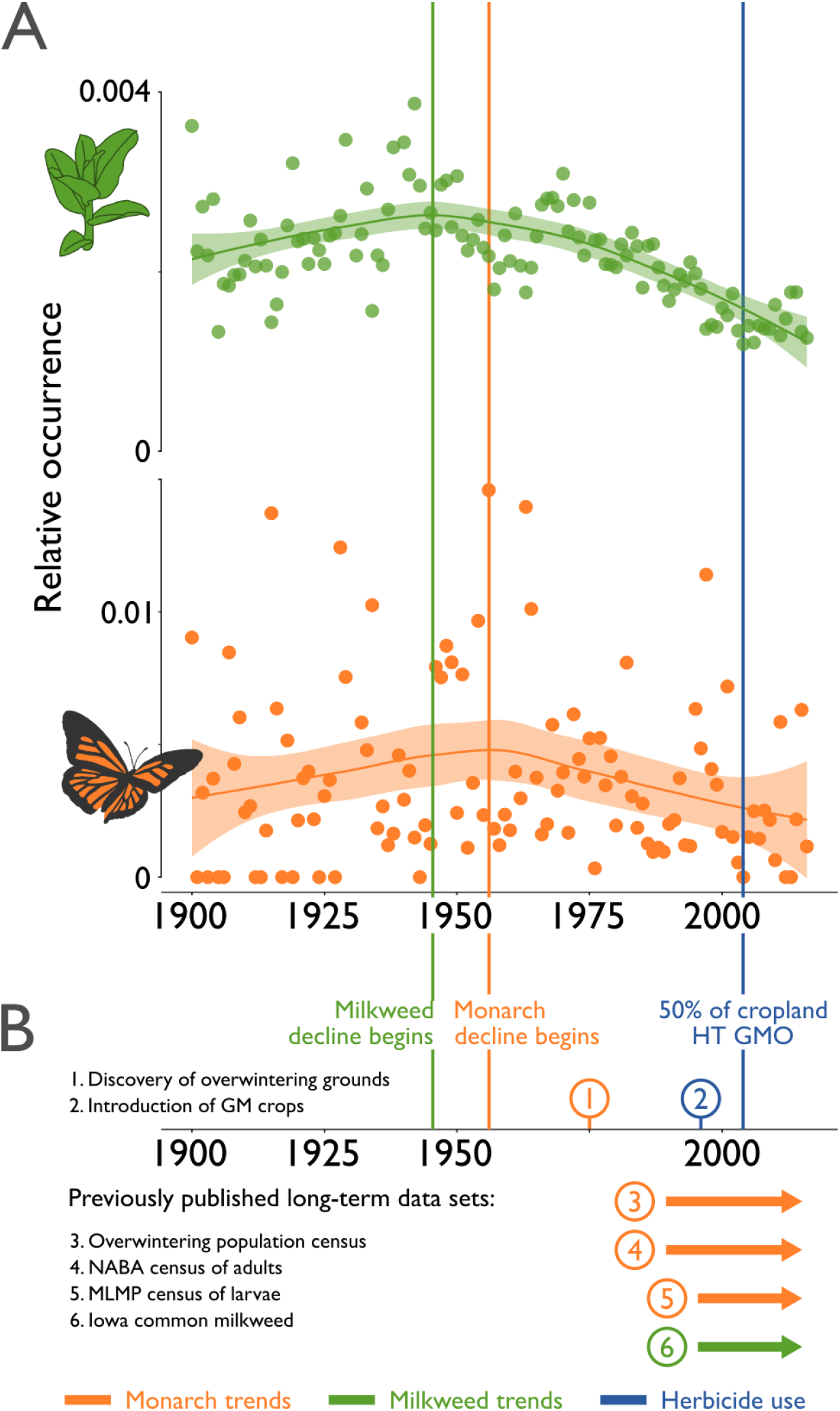
Museum specimens reveal long-term trends in monarchs and milkweed. **1A**: green points show annual occurrences for milkweed spp.; orange points for monarchs; lines and shading indicate smoothed mean and 95% confidence intervals, calculated using the Loess smoothing method implemented in ggplot2^19^, with the default smoothing span. Green and orange vertical lines indicate the approximate beginning of the decline for milkweed and monarchs, respectively. The blue vertical line indicates the point at which half of all corn, soybeans, and cotton are GM varieties. **1B:** indicates (1) the discovery of the monarch overwintering grounds in Mexico; (2) the introduction of GM crops; (3) the winter population census at the Mexican overwintering grounds^20^; (4) the summer NABA census of adults (available from^6^); (5) the summer MLMP census of eggs and larvae (available from^12^); (6) the summer census of Iowa *A. syriaca* abundance (available from^12^).

**Table 1:** Occurrence records used in this study

| Data source | *Asclepias* records | Total plant records |
| --- | --- | --- |
| Global Biodiversity Information Facility | 14,458 | 1,264,715 |
| Consortium of Midwest Herbaria | 21,458 | 3,208,245 |
| Online Virtual Flora of Wisconsin | 2496 | 362,789 |
| Minnesota Biodiversity Atlas | 1098 | 144,254 |
| Total | 39,510 | 4,980,003 |

|  | *D. plexippus* records | Total lepidoptera records |
| --- | --- | --- |
| Symbiota Collections of Arthropods Network | 1191 | 323,611 |

For both monarchs and milkweed, relative occurrence shows substantial year-to-year variation, but the trend over the twentieth century is nevertheless clear. Monarch occurrence is linked to milkweed occurrence: both species increase early in the century, milkweeds peak slightly before monarchs (around 1945, and 1955, respectively), and both suffer a two-fold decline between then and 2016.

The monarch data show substantially more year-to-year variation than the milkweed data, likely both because the milkweed data is based on nearly 30 times as many records, and because the annual variation in monarch populations is considerable: for instance, the overwintering population size commonly experiences two-to-fivefold changes from year to year^11^.

### Comparison of our trends from museum specimens to other data sets

Year-to-year variation in both monarchs and milkweed obscured the correlation between them (r = 0.16, *p* = 0.08), but our estimates of occurrence correlate with other monarch butterfly datasets over the approximately 20-year period over which they overlap (data from^6, 12,20^). In particular, our monarch occurrences correlated with estimates of egg abundance from the Monarch Larvae Monitoring Project (MLMP) (r = 0.65, *p* < 0.01). They likewise correlate, though not statistically significantly, with the North American Butterfly Association (NABA) estimates of adult abundances, once corrected for land cover using the method of^12^ (r = 0.48, *p* = 0.06). The correlation between milkweed abundance and the size of the Mexican overwintering population the following winter is not statistically significant (r = 0.34, *p* = 0.14). To further investigate this relationship, we restricted our analysis to those states from which the bulk of the overwintering monarch originate^21^. Milkweed abundance in this core area does predict the size of the subsequent overwintering monarch population (r = 0.45, *p* = 0.04). Further comparisons, alongside plots of the relationships between the different estimates of monarch and milkweed abundances are provided in the Supporting Information.

### *Occurrence trends in individual* Asclepias *species from 1900-2016*

Particular species may be more or less likely to be collected as herbarium specimens^22^; therefore, changing communities within the genus could bias the genus-wide trends in *Asclepias*. To exclude this possibility, we also looked at the population trends for the 10 most abundant individual species, as the particular collection biases for each species should be relatively constant over time.

We found that each of the ten most abundant milkweed species showed a trend similar to the genus-wide trend (Figure 2). In particular, all ten species showed a marked decline toward the end of the twentieth century. The beginning of this decline was species specific. For instance, *A. speciosa* and *A. viridis* decline over most of the studied period; *A. amplexicaulis* and *A. asperula* have brief periods of increase before a decline beginning around the 1920s, while the declines for *A. incarnata* and *A. verticillata* begin much later, around the 1970s.

**Figure 2:**
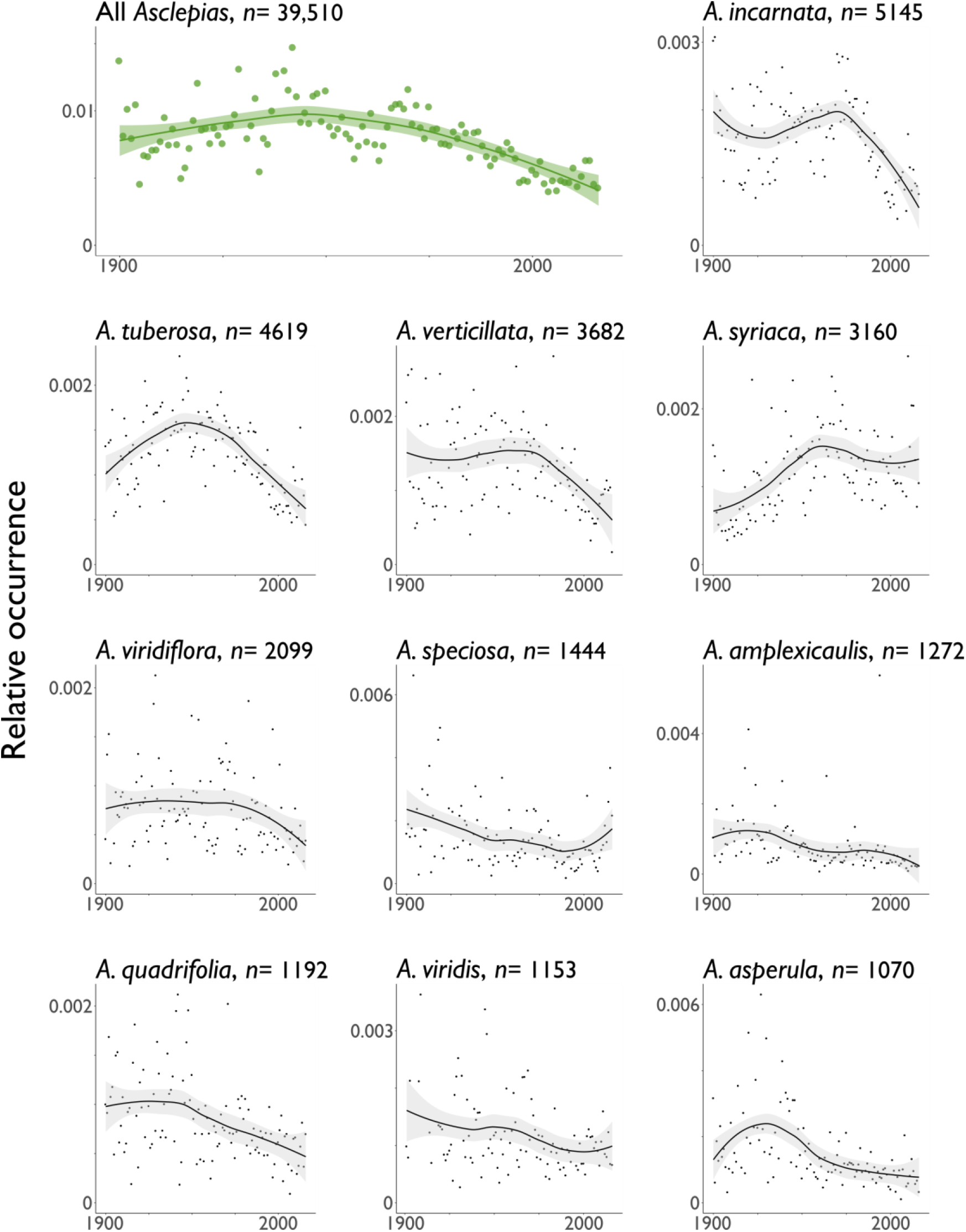
Genus-level milkweed decline over the twentieth century is recapitulated in the ten most common *Asclepias* species. Invasions of the continental United States are apparent from museum specimens. The total number of specimens collected is shown next to each species. Points indicate occurrence for each year, lines and shading indicate smoothed mean and 95% confidence intervals. Smoothing was done using the Loess smoothing method implemented in ggplot2^19^, with the default smoothing span. Because different species have different ranges, the occurrences for each species do not add up to the occurrence for the genus as a whole.

While each species declines over the latter part of the twentieth century, these declines are relatively slow for *A. syriaca* and *A. speciosa*, and in fact both species show signs of slight increase in population size after 2000. This may be an artefact of noise in the data, as there are fewer records digitized after 2000 (Figure S1). However, it may also show increases in milkweed due to monarch conservation efforts that encourage the planting of milkweed (e.g.^23, 24^).

The relatively slow declines in these two species mean that these species now account for a greater proportion of the total milkweed records than they did at the beginning of our study period. This change in the makeup of the community of milkweeds is visualized in Figure 3.

**Figure 3:**
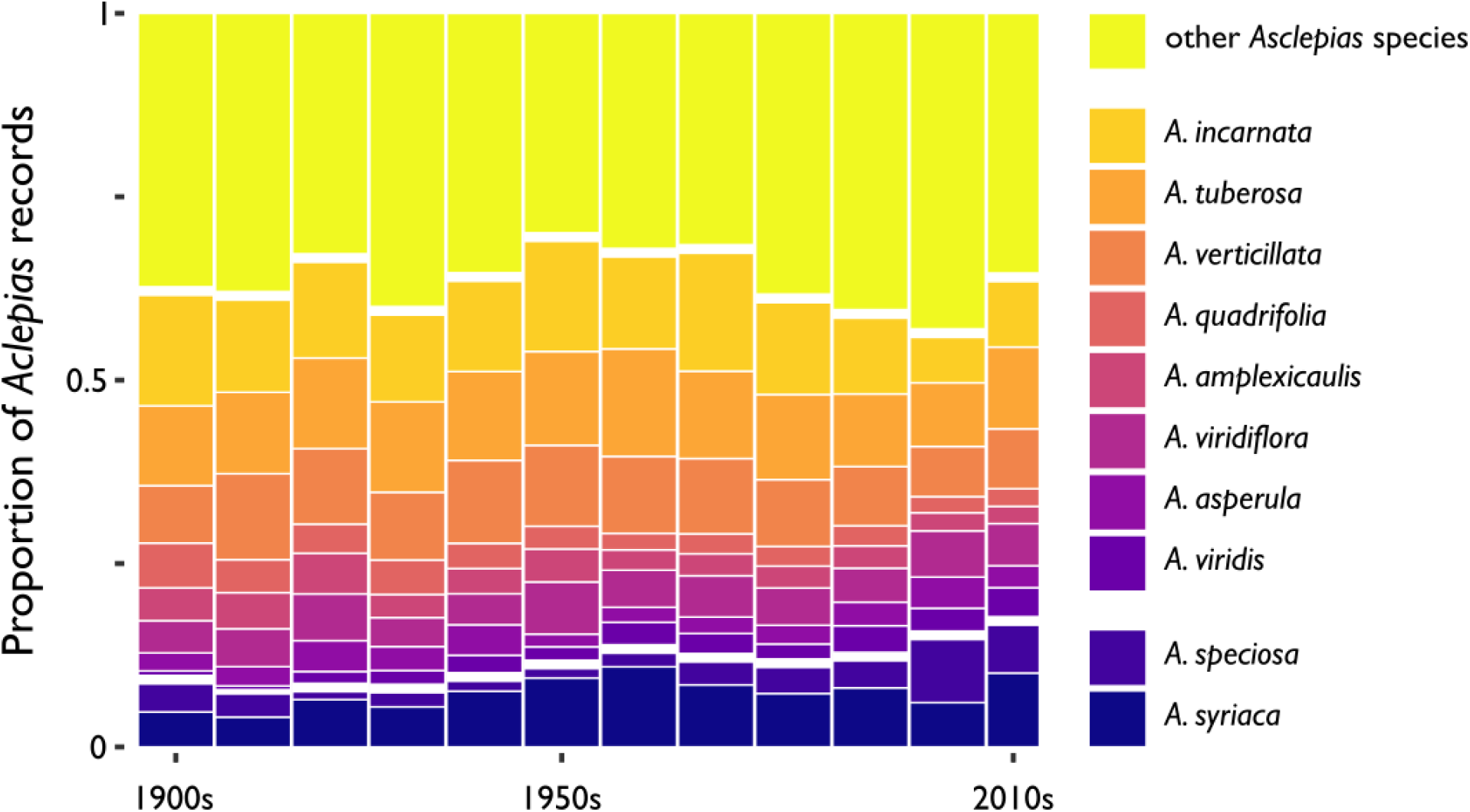
Changes in composition of *Asclepias* records over time. *A. syriaca* and *A. speciosa* make up a greater proportion of *Asclepias* records in the past few decades than early in the twentieth century.

### Multimodel inference

To investigate which changes in farming practice predict monarch and milkweed decline, we focus on the primary host plant of the monarch, *A. syriaca*, the common milkweed. Over the period of 1950-2006, for which we have good data on United States agricultural practices, *A. syriaca* increased in occurrence from 1950-1970, then subsequently declined (Figure S8). We performed multi-model inference to test whether changing agricultural practices in the United States (Figure S8) predict *A. syriaca* abundance. In every selected model, *A. syriaca* abundance was negatively correlated with number of farms, which declined over the period 1950-2006 as smaller farms consolidated. Most selected models included at least one other factor: total area of farmland, fertilizer use, and glyphosate use all appeared in at least one selected model, although their effect sizes were much smaller than “number of farms”, and confidence intervals overlapped zero for these three traits, leaving open the possibility that they do not predict *A. syriaca* abundance.

## Discussion

Both monarchs and milkweeds increase over of the beginning of our study period before declining to their present abundance. The increase of both groups in the early 1900s is interesting, as some authors suggest that milkweeds and monarchs experienced a range expansion in the late 1800s, driven by the conversion of eastern forests to farmlands^10, 25^. Our early-twentieth-century increases in monarch and milkweed may reflect the tail end of such a trend, although the number of records at the beginning of the century is probably too small to be certain about the degree or precise timing of such an increase.

The decline for both monarchs and milkweed appears monotonic, suggesting that the well-studied decline from 1993-date is part of a larger trend beginning in the middle of the last century. While the monarch trend closely follows the milkweed trend, our data is correlational, and thus it is difficult to distinguish between several competing hypotheses.

It could be the case that the declines in milkweed cause monarch declines (the “milkweed limitation hypothesis” of, e.g.,^2, 12^), or monarch declines may be caused by some other factor which is correlated with milkweed declines, such as loss of nectar resources for the adults (^6, 26^) or severe weather and changing climate (^2, 27^), or more than one of the above.

When looking at the declines for each individual *Asclepias* species, these declines were the least marked in *A. syriaca* and *A. speciosa* (Figures 2 and 3). These two species are the hostplants of the majority of monarch larvae in the central and northern United States^15, 28,29^. However, Martin and Lynch^29^ and Brower^10^ both hypothesized that monarchs’ reliance on *A. syriaca* and *A. speciosa* is a recent phenomenon, caused by the (relative) increase of these disturbance-loving species on a human-disturbed landscape. Our results are the first quantitative evidence that *A. syriaca* and *A. speciosa* have recently increased their share of the *Asclepias* community: these two species account for an increasing proportion of the total *Asclepias* records over time, almost doubling from 9% in 1900-1909 to 17% in 2010-2016 (Figure 3). *A. syriaca* and *A. speciosa* are relatively poorly chemically defended^29^; since monarchs get their chemical defenses from their larval hostplants, current populations of monarchs may possess worse chemical defenses than monarchs did 50-100 years ago, as suggested by^10, 29^.

Based on our multi-model inference results, we suggest a preliminary hypothesis to explain the rise and fall of *A. syriaca* abundance after 1950: Over the course of the 1950s and 1960s, many small farms rapidly consolidated into fewer, larger farms (Figure S8). We suggest this likely reduced the area of uncultivated divisions between different properties, benefitting *A. syriaca*, which thrived in the relatively competitor free areas between crop rows in the fields themselves^30^. However, beginning in the 1970s, the rate of farm consolidation greatly slowed, and this was no longer enough to buoy *A. syriaca* populations against threats such as a decline in total area of farms, or increasing use of glyphosate (and other herbicides) in the fields themselves. Our global model explains about 18% of the variation of common milkweed, leaving much to be done in explaining its changing occurrence patterns, especially outside of agricultural land.

Herbicide-resistant crops have been identified as a major cause of declines of both monarch and milkweeds, a conclusion that has impacted many people’s opinions of GM crops. Our results show that the well-studied decline in monarch populations from 1993-date is part of a much larger trend, with monarch declines beginning in the 1950s and continuing steadily until the present day, following a decline in milkweed host plants. These results clearly indicate that herbicide resistant crops are not the only culprit, as declines in both monarchs and milkweed begin well before GM crops were introduced. Solely focusing on GM crops at the expense of other potential drivers will hinder our ability to address and reverse the worrying declines in these species.

## Materials and Methods

### Plant and insect records

We gathered online herbaria records of preserved plant specimens from four sources: the Global Biodiversity Information Facility^31^, the Consortium of Midwest Herbaria^32^, the University of Minnesota Bell Museum of Natural History^33^, and the Online Virtual Flora of Wisconsin^34^ (Table 1). Records were cleaned to include only tracheophytes (i.e., all vascular plants) collected in the contiguous United States between 1900 and 2016, inclusively. Records from several institutions were found in more than one of our data sources: in these cases, the duplicated records were deleted from all but one of the data sources. We also gathered online museum records of insects from the Symbiota Collections of Arthropods Network^35^ (Table 1). Records were cleaned to include only preserved specimens of lepidoptera (i.e., butterflies and moths) collected in the same time period and location as the plant records. Data cleaning and all statistical analyses described below were done in R version 3.4.2^36^, and the scripts used are available on Dryad.

For each record, we used the associated latitude and longitude to estimate the kind of landscape upon which that specimen was collected. To do this, we consulted the USDA National Agricultural Statistics Service Cropland Data Layer^37^. These data provide estimates of land cover for the continental United States since 2008. We used these data to categorize specimens as being collected from one of the following categories: developed land (Cropland Data Layer categories 82, 121-124), crop land (CDL categories 1-60, 66-77, 204-254), natural land (CDL categories 63-65, 87, 88, 112, 131-152, 190, 195), grassland (CDL categories 61, 62, 171, 176, 181; this includes both agricultural grasslands such as fallow fields and non-agricultural grasslands), or water (CDL categories 83 and 111). Some points changed cover category over the period since 2008. If land cover information was available for the year in which a specimen was collected, we assigned that specimen that cover category. Otherwise, if one cover category was found at that point during more years than any other cover category, we assigned that specimen that plurality cover category. If there was no plurality cover category, we did not assign a cover category.

Since land cover data only goes back to 2008, these assignments should be viewed as preliminary, as many specimen collection sites no doubt changed land cover between when that specimen was collected and 2008. For instance, some specimens categorized as being collected from “developed” land were likely collected from pasture or agricultural land which has since urbanized.

Monarch butterflies in the United States are divided into two distinct migratory populations: east and west. In this study, we focus on the eastern population of butterflies and their host plants, and so we excluded records from states west of the continental divide (Washington, Oregon, Colorado, Idaho, Nevada, Utah, Arizona) from our data set.

### *Occurrence trends in the genus* Asclepias *from 1900-2016*

Our cleaned, eastern data set included 39,510 records of *Asclepias* species. However, raw number of *Asclepias* specimens is a poor metric of occurrence, because collection effort has varied over the course of our study period (Figure S1). In order to control for varying collection effort from year to year, we calculated relative occurrence of *Asclepias* by taking the quotient of the number of *Asclepias* records collected each year divided by the total number of tracheophyte records collected within the range of *Asclepias* species. We estimated the genus-wide range of *Asclepias* with a bounding box containing all *Asclepias* records except for the most extreme 1% in each direction, north, south, east, and west. We did this analysis within each of the four data sources, then combined the four by calculating the average, weighting each data set by the number of *Asclepias* records in that data set in that year. When visualizing the trends for *Asclepias*, we removed some years which were substantial outliers (1930 and 1939), i.e., falling greater than three standard deviations away from the mean annual occurrence.

To confirm the sensitivity of this relative occurrence metric to real changes in population size, we followed a similar procedure for several other herbaceous plants, which are known to have invaded the United States during the time period of this study. As described in the Supporting Information, these invasive species showed marked increases in their relative occurrence over the twentieth century (Figure S5).

Shorter-term trends in milkweed decline appear to vary by land cover category; e.g., declines in crop fields land may be much steeper than declines in non-agricultural land, like roadsides^38^. We investigated whether this was the case for our long-term trends. As described in the Supporting Information, *Asclepias* showed a decline in occurrence across all studied land cover types (Figure S6).

### *Occurrence trends in individual* Asclepias *species from 1900-2016*

Particular species may be more or less likely to be collected as herbarium specimens^22^: for instance, an *Asclepias* species that is commonly found near road sides may be collected more commonly than a second *Asclepias* species that is equally common, but found in less convenient locations. Therefore, any decreasing trends in *Asclepias* occurrence could have two explanations: first, that there were fewer *Asclepias* plants found over time; second, that the total number of *Asclepias* plants remained constant, but that the more easily collected species declined while the less easily collected species increased, leading to an apparent decline when looking at all *Asclepias* records at once.

To test between these two possibilities, we also looked at the population trends for individual species, as the particular collection biases for each species should be relatively constant over time. We looked at individual trends for the 10 most abundant milkweed species in our data set: *A. incarnata, A. tuberosa, A. verticillata, A. syriaca, A. viridiflora, A. speciosa, A. amplexicaulis, A. viridis, A. quadrifolia*, and *A. asperula.* Records of these 10 species combined made up 63% of the total data set. For each species, we calculated its range and occurrence as described above for the *Asclepias* genus. When visualizing the trends for individual species, we removed some years which were substantial outliers, i.e., falling greater than three standard deviations away from the mean annual occurrence (A. *amplexicaulis:* 1903, 1909, 1988; *A. asperula:* 1936, 1940; *A. quadrifolia:* 1924; *A. speciosa:* 1943, 1970; *A. tuberosa:* 1937; *A. verticillata*, 1921; *A. viridiflora;* 1904; *A. viridis*, 1904, 1918, 1994). We also divided the *Asclepias* records into 10-year bins, and calculated the relative proportion of each species over time (we did not identify or remove outliers in this part of the analysis).

### *Occurrence trends for* Danaus plexippus *from 1900-2016*

Our cleaned, eastern data set included 1191 records of *Danaus plexippus.* As with *Asclepias*, we calculated the geographic range of *D. plexippus* as described above, then estimated the occurrence of *D. plexippus* for each year by comparing the number of *D. plexippus* specimens collected to the total number of lepidoptera specimens collected within the *D. plexippus* range. When visualizing the trends for *D. plexippus*, we removed some years which were substantial outliers (1930 and 1931), i.e., falling greater than three standard deviations away from the mean annual occurrence.

### Comparison of our trends from museum specimens to other data sets

Using Pearson’s correlation coefficient, we compared the occurrence of milkweeds and monarchs from our museum data both to each other, and also to estimates of monarch and milkweed abundance from other datasets. We examined three other data sets: estimates of the size of the monarch overwintering population from 1994-2014^20^, Monarch Larva Monitoring Project (MLMP) estimates of immature (egg stage) monarch population sizes in the summer breeding grounds from 1999-2014^12^, and North American Butterfly Association (NABA) estimates of adult monarch population sizes in the summer breeding grounds from 1993-2014^6^. For the latter two data sets, we also employed the corrections for changes in land cover described by Pleasants *et al*.^12^.

A relatively small number of states contribute disproportionately to the Mexican overwintering population^21^. Therefore, we also calculated milkweed occurrence in these states alone, using the methods described above, but including only records from Texas, Oklahoma, Missouri, Illinois, Indiana, and Ohio. We compared these estimates of milkweed occurrence from the core area to the size of the overwintering population.

### Agricultural data

We gathered data on selected agricultural practices in the United States, namely, the number of farms and other agricultural operations such as ranches and tree nurseries^39^, the total area of farmland^40^, the amount of nitrogen and phosphorus fertilizers used^41, 42^, and the amount of glyphosate herbicide used^43^. Data for all of these were available between 1950 and 2006, inclusive. Nitrogen and phosphorus fertilizer use were highly correlated with each other (Figure S2), and so we combined them into a single variable by scaling both variables to have a mean of zero and a standard deviation of 1, then adding the scaled variables to produce a metric of total nitrogen-plus-phosphorus fertilizer used.

Data on glyphosate use were only available at the national level; data on the other three factors, however, were available at the state level. We divided the states into six regions (Figure S3) with relatively homogenous agricultural practices, combining the data for each state. We divided the *A. syriaca* and tracheophyte records gathered above by region, then used these to calculate the relative occurrence of *A. syriaca* within each region.

Since there was some degree of variation from year to year, we pooled the regional data into five year bins. As the year-to-year data for *A. syriaca* contained several outlying data points, we took the median value for *A. syriaca* occurrence in each five year bin, as this greatly lessened the ability of outlier data points to effect the model compared to year-to-year data or taking the median of the five year period. Using the median rather than mean or single-year bins accordingly increased the predictive power of the global model and increased our ability to distinguish different models using AIC (results not shown). We averaged the agricultural data across the five year period (or the two year period, in the case of the 2005-2006 bin). The agricultural data was then standardized so that within each factor, the mean was zero and the standard deviation was one.

### Multimodel inference

To test which changes in agricultural practice had an effect on *A. syriaca* abundance, we performed multimodel inference using the MuMIn package^44^. We ran 16 different mixed-effects models, each with *A. syriaca* abundance as the response variable. The fixed-effects variables were some combination of total area of farmland, number of farms, fertilizer used, and glyphosate herbicide used, and a random-effect of geographic region on the y-intercept was also used. The 16 models include every possible combination of the four fixed-effect variables, including a null model with only an intercept term.

We calculated the relative quality of each model using AIC, retaining in our analysis any model within 4 AIC units of the highest-quality model. The effect of each variable on *A. syriaca* abundance was averaged across all the retained models, weighting by the relative likelihood of each model. When a variable was not found in a model, it was considered to have an effect of zero. The results of this analysis are shown in Table 2 and discussed in the main text.

**Table 2:**
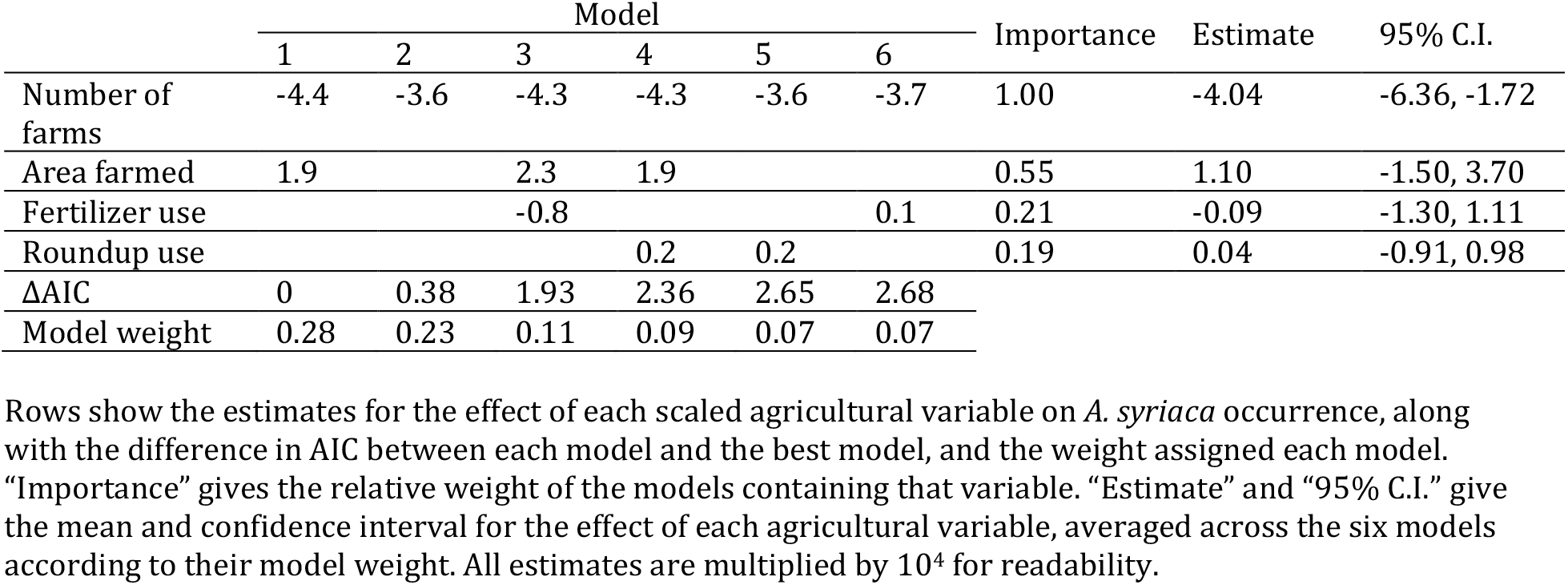
Results of multi-model inference

|  | Model |  |  |  |  |  | Importance | Estimate | 95% C.I. |
| --- | --- | --- | --- | --- | --- | --- | --- | --- | --- |
|  | 1 | 2 | 3 | 4 | 5 | 6 |  |  |  |
| Number of farms | -4.4 | -3.6 | -4.3 | -4.3 | -3.6 | -3.7 | 1.00 | -4.04 | -6.36, -1.72 |
| Area farmed | 1.9 |  | 2.3 | 1.9 |  |  | 0.55 | 1.10 | -1.50, 3.70 |
| Fertilizer use |  |  | -0.8 |  |  | 0.1 | 0.21 | -0.09 | -1.30, 1.11 |
| Roundup use |  |  |  | 0.2 | 0.2 |  | 0.19 | 0.04 | -0.91, 0.98 |
| ΔAIC | 0 | 0.38 | 1.93 | 2.36 | 2.65 | 2.68 | | | |
| Model weight | 0.28 | 0.23 | 0.11 | 0.09 | 0.07 | 0.07 |  |  |  |

Rows show the estimates for the effect of each scaled agricultural variable on *A. syriaca* occurrence, along with the difference in AIC between each model and the best model, and the weight assigned each model. “Importance” gives the relative weight of the models containing that variable. “Estimate” and “95% C.I.” give the mean and confidence interval for the effect of each agricultural variable, averaged across the six models according to their model weight. All estimates are multiplied by $10^4$ for readability.

For each plant data source, we downloaded all available records as of March 2018. For insects, we downloaded all records matching a higher taxonomy search for “Lepidoptera” as of April 2018. In both cases, we then filtered records to produce these sample sizes.

## Data availability

Data tables used for all analyses, along with R scripts for each analysis, will be made available on dryad.

## Acknowledgements

The insect collections staff at the University of Minnesota and University of Kansas aided us by providing *D. plexippus* records through the Symbiota Collections of Arthropods Network. We are grateful to DS De La Mater and the Milkweed Research Group at William & Mary for their feedback on the manuscript. JHB was supported by a postdoctoral fellowship in the William & Mary Environmental Science and Policy Program funded by the Andrew W. Mellon Foundation.

## Author Contributions

JHB, HJD, and JRP designed the study. JHB gathered the data and performed statistical analyses. JHB wrote the manuscript with input from HJD and JRP.

## Supporting Information

**Figure S1:**
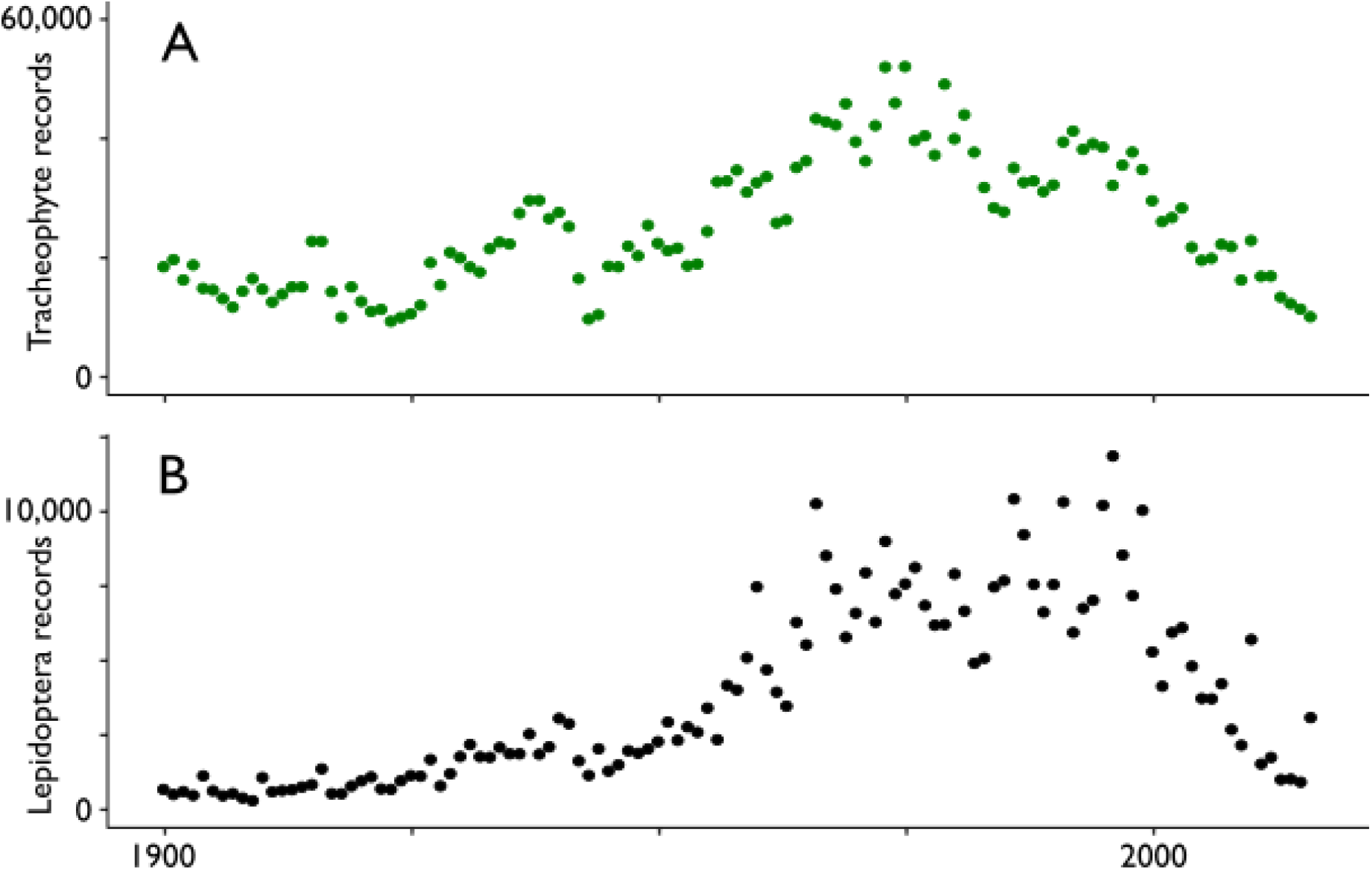
Collection effort varies over time. A) shows the total number of cleaned tracheophyte records from eastern states for each year; B) shows the same for lepidoptera records.

**Figure S2:**
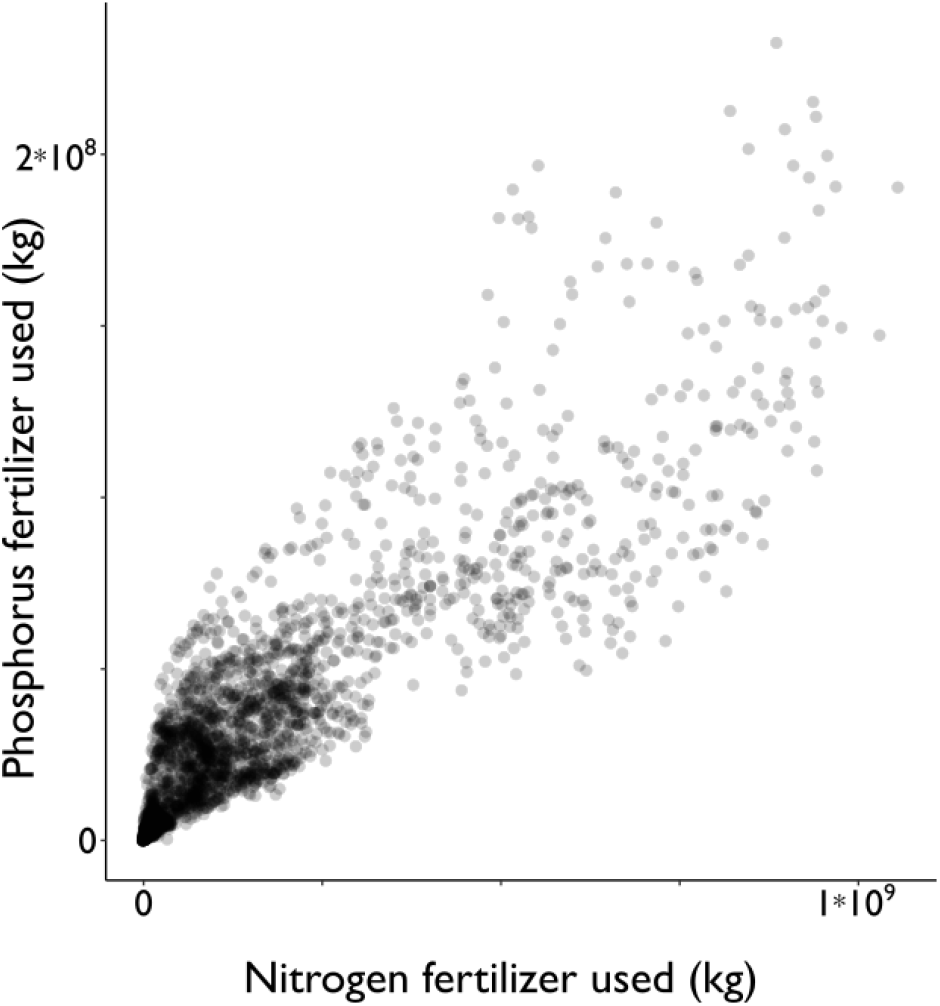
Nitrogen and phosphorus use are tightly correlated. (r = 0.88, p << 0.001). Each point represents the fertilizer use for a single state (include all 48 states in the continental United States) in a single year from 1950-2006.

**Figure S3:**
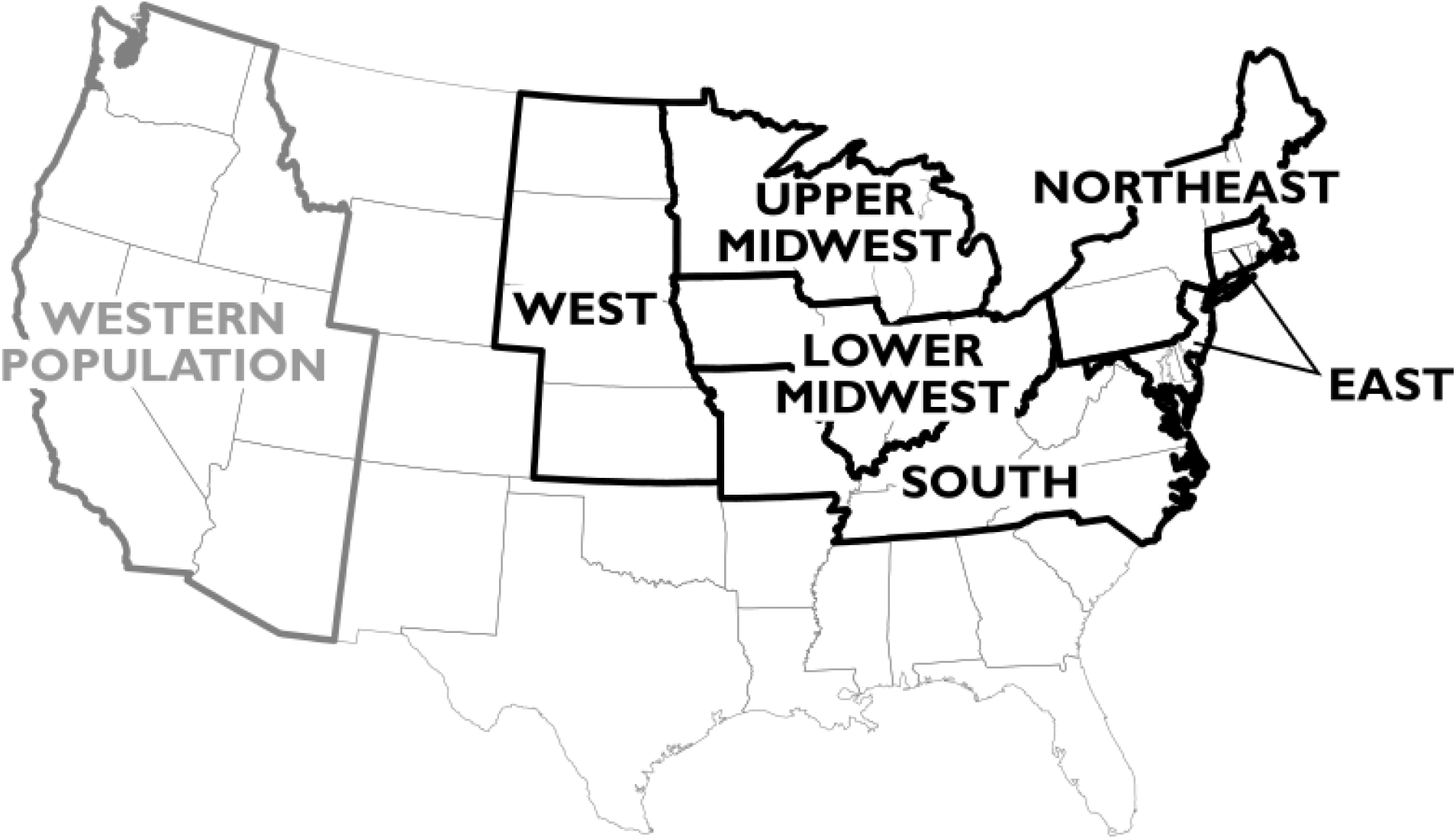
The six agricultural regions used in the *A. syriaca* model. The six regions are outlined in black. States how to the western population of *D. plexippus* are also indicated.

### Use of genetically modified crops in the United States

The main genetically modified (GM) crops in the United States are corn, soybeans, and cotton (National Academies of Science, Engineering, and Medicine 2016). To estimate the total area of US cropland dedicated to GM crops, we gathered estimates of what proportion of each of these crops were genetically modified to express resistance to herbicide. Our estimates came from four sources: the USDA-NASS report *Acreage* (1995-2016) for the period 2000-2016; Fernandez-Cornejo and McBride (2002) for cotton and soybeans for the period 1996-1999; Fernandez-Cornejo and McBride (2000) for corn 1996-1997; the USDA-NASS report *Crop Production* (1999) for corn 1998-1999. We multiplied these to the total area planted for each of these three crops (USDA-NASS 1995-2016) to estimate the total number of acreages planted with GM crops, which we compared both to the total acreage planted for corn, soybeans, and cotton, and to the total acreage planted for all crops (USDA-NASS 1995-2016).

The prevalence of genetically modified herbicide resistant crops increased steadily since their introduction in 1996 as shown in Figure S4.

**Figure S4:**
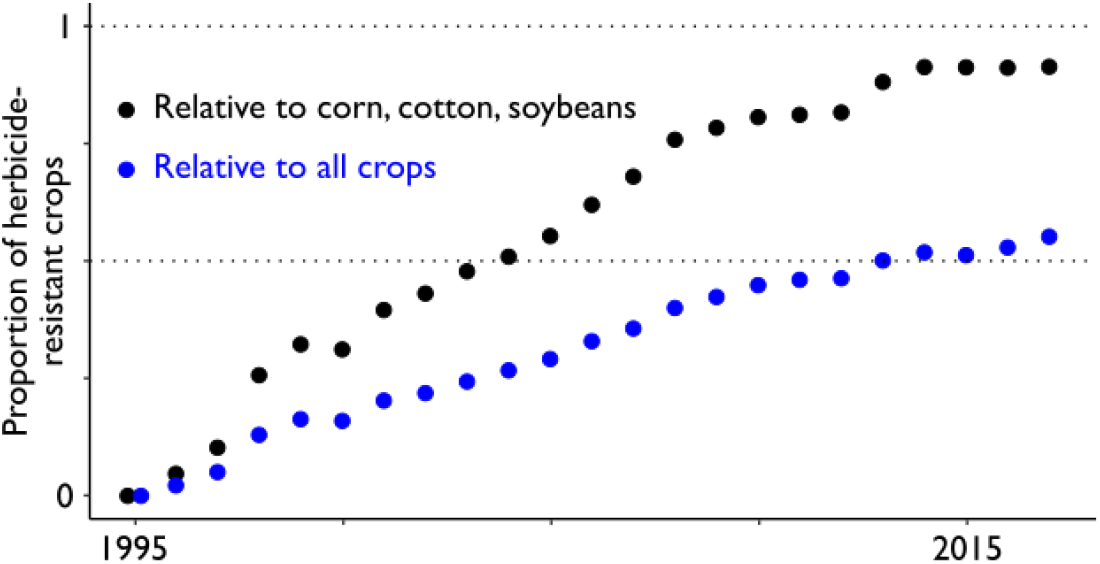
Increase in herbicide-resistant (HR) GM crops in the United States. HR crops were introduced in 1996. Each point shows the proportion of HR GM acreage of corn, cotton, and soybeans as a proportion of all corn, cotton and soybeans acreage (black points), and as a proportion of all crops (blue points). Half of all corn, cotton, and soybeans were HR by 2004; half of all crops by 2013.

### Occurrence trends in other plant species from 1900-2016

To confirm the sensitivity of this analysis to real changes in population size, we did a similar procedure for several species with ongoing invasions of the United States during the time period of this study: garlic mustard, *Alliaria petiolata* (Nuzzo 1993); purple loosestrife, *Lythrum salicaria* (Stuckey 1980); Japanese stiltgrass, *Microstegium vimineum* (Hunt and Zaremba 1992); and kudzu, *Pueraria montana* (Forseth and Innis 2004). *P. montana* is a synonym with *P. lobata*, and some data sets had records for both species names; in this case, we combined *P. lobata* and *P. montana* records. For each invasive species, we compared the total number of records for that species to the total number of tracheophyte records collected within that species’ range. Species’ ranges were calculated as described for *Asclepias* in the main text.

When visualizing the trends for individual species, we removed some years which were substantial outliers, i.e., falling greater than three standard deviations away from the mean annual occurrence *(Alliaria petiolata:* 2002; *L. salicaria:* 2009, 2016; *M. vimineum:* 2001, 2004; *P. montana:* 1966, 1967).

In each case, we detected marked increases in occurrence over the course of the twentieth century for these plants known to be invasive in the United States over that period (Figure S5).

**Figure S5:**
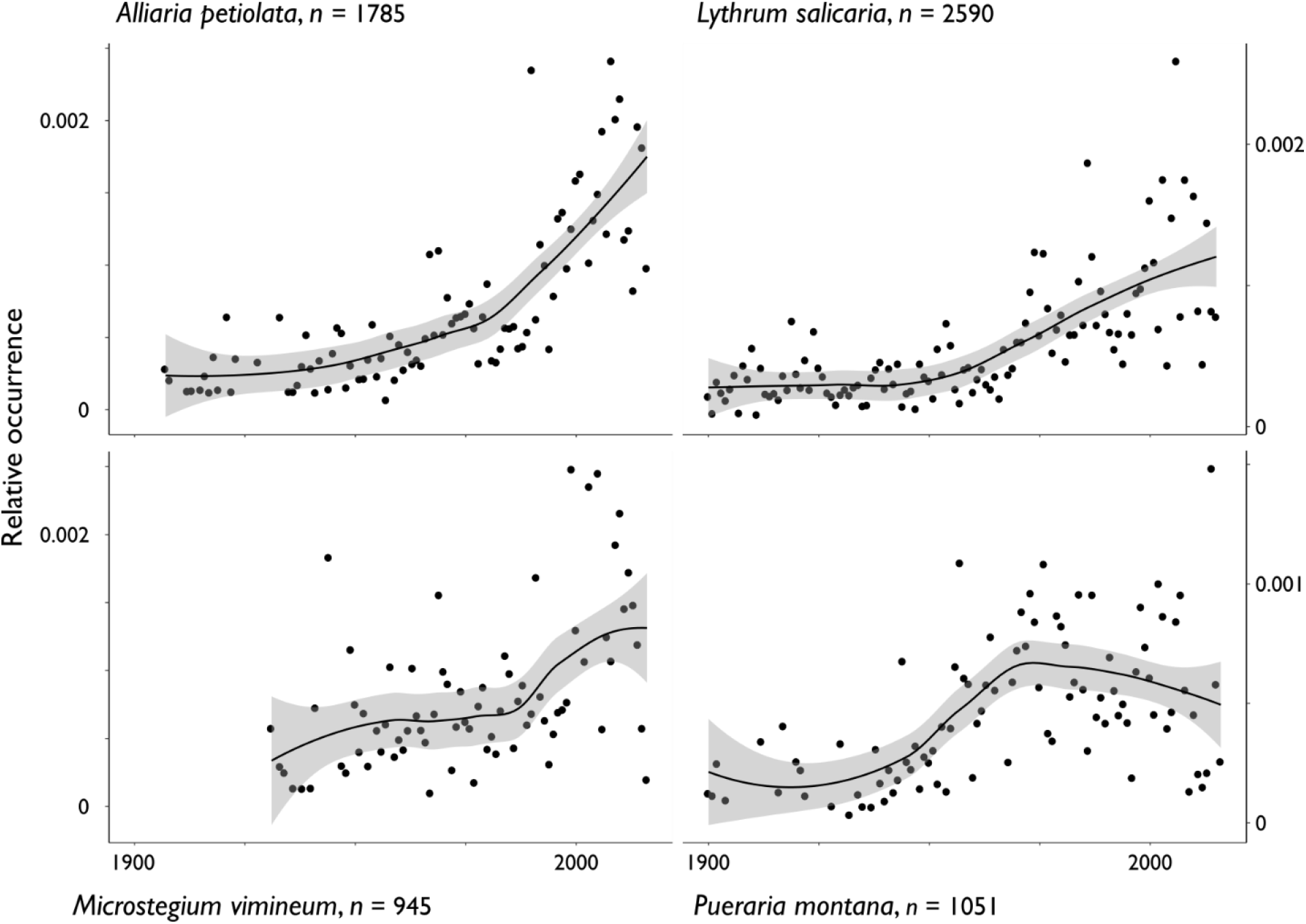
Invasions of the continental United States are apparent from museum specimens. The total number of specimens collected is shown next to each species. Points indicate occurrence for each year, lines and shading indicate smoothed mean and 95% confidence intervals. Smoothing was done using the Loess smoothing method implemented in ggplot2 (Wickham 2009), with the default smoothing span.

### Breaking down occurrence trends by land cover category

Shorter-term trends in milkweed decline appear to vary by land cover category; e.g., declines in crop fields land may be much steeper than declines in non-agricultural land, like roadsides (Hartzler 2010). We investigated whether this was the case for our long-term trends.

To calculate occurrence on each land cover category, we compared the number of *Asclepias* specimens assigned to that category to the total number of specimens (of all categories) collected in that year, as above. We did not do this for *D. plexippus*, for which there were not enough records to subdivide. To account for the fact that the number of specimens assigned to a land cover category changes over time (as more specimens are associated with geographic coordinates), we divided this occurrence by the proportion of *Asclepias* specimens collected that year which were assigned a land cover category. Finally, we averaged each data source (i.e., GBIF, WIS, etc) separately, weighting them the same as described in the main text methods section, “Occurrence trends in the genus Asclepias from 1900-2016”. When visualizing the trends for each land cover category, we removed some years which were substantial outliers, i.e., falling greater than three standard deviations away from the mean annual occurrence (records from crops: 1971, 1975, 1983; developed land: 1900; grassland: 1929, 1967; natural land: 1939).

We found declines in milkweed occurrence in all four categories of land cover (Figure S6). In the case of cropland, grassland, and natural land, we saw an increase in the early twentieth century that predated the decline in the second half of the century. In the case of developed land, we saw a steady decline, although this could be because many sites that are currently developed were in fact in another land cover category before urbanization. Thus, the count of records from developed land is likely inflated in the early part of the twentieth century.

**Figure S6:**
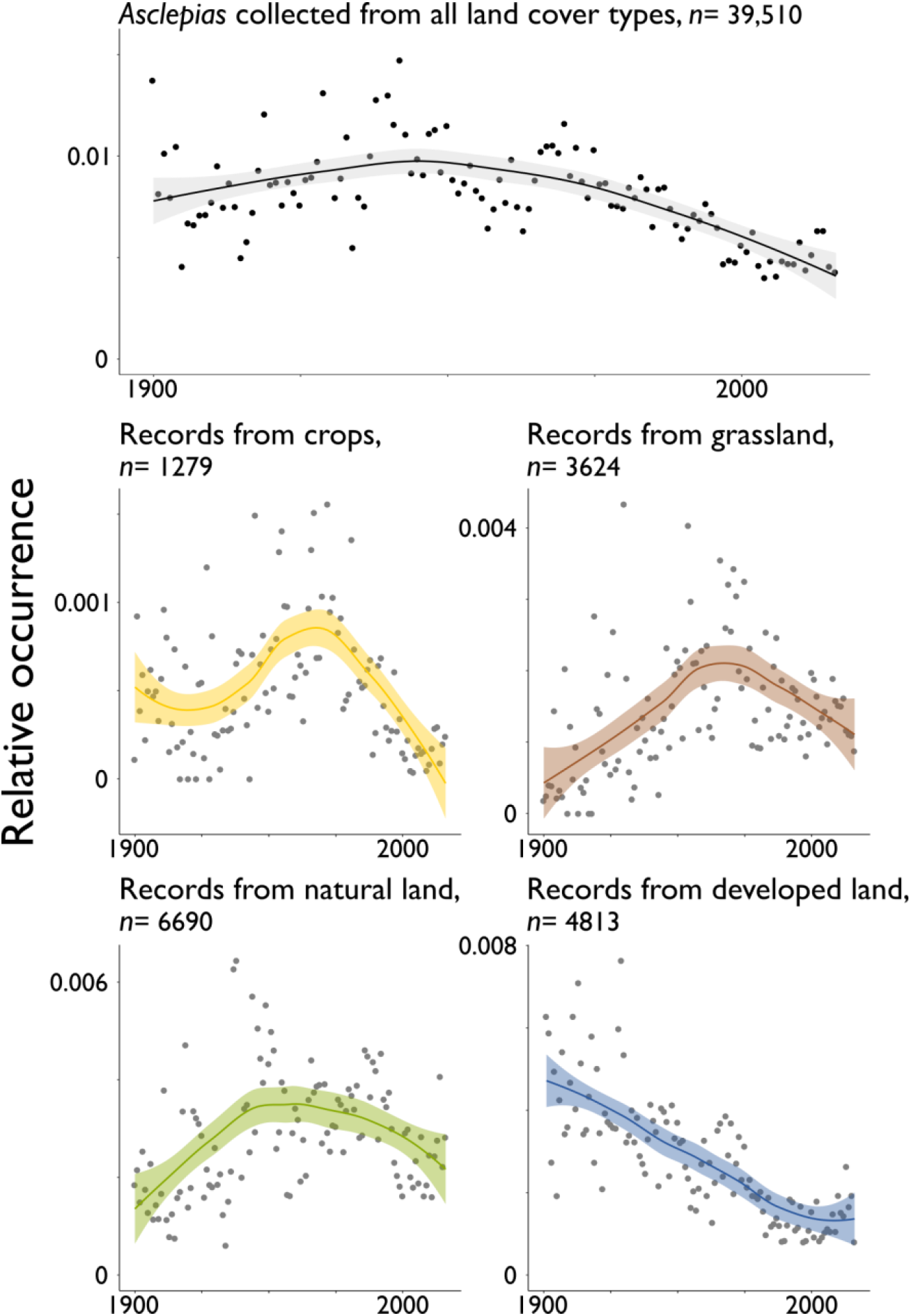
Milkweed decline over the twentieth century is seen across land use types. The total number of specimens collected on that land cover type is shown above each graph. Points indicate occurrence for each year, lines and shading indicate smoothed mean and 95% confidence intervals. Smoothing was done using the Loess smoothing method implemented in ggplot2 (Wickham 2009), with the default smoothing span. Because land cover was not determined for all records, the occurrences for each land cover type do not add up to the occurrence for the genus as a whole.

Using Pearson’s correlation coefficient, we compared the occurrence of milkweeds and monarchs from our museum data both to each other, and also to estimates of monarch and milkweed abundance from other datasets. We examined three other data sets: estimates of the size of the monarch overwintering population from 1994-2014 (Monarch Watch 2016), Monarch Larva Monitoring Project (MLMP) estimates of immature (egg stage) monarch population sizes in the summer breeding grounds from 1999-2014 (Pleasants *et al.* 2017), and North American Butterfly Association (NABA) estimates of adult monarch population sizes in the summer breeding grounds from 1993-2014 (Inamine *et al.* 2016). For the latter two data sets, we also employed the corrections for changes in land cover described by Pleasants *et al.* (2017).

A relatively small number of states contribute disproportionately to the Mexican overwintering population (Flockhart *et al.* 2013). Therefore, we also calculated milkweed occurrence in these states alone, using the methods described above, but including only records from Texas, Oklahoma, Missouri, Illinois, Indiana, and Ohio. We compared these estimates of milkweed occurrence from the core area to the size of the overwintering population.

We found no correlation between our estimate of milkweed abundance with our estimate of monarch abundance. However, there was substantial year-to-year variation, which may have obscured the overall trend. To test this hypothesis, we grouped monarch and milkweed occurrence into five-year bins, taking the median occurrence for each bin, and measured the correlation between these two data sets.

There is little correlation between our *D. plexippus* occurrence and NABA citizen-science counts of adult butterflies (r = 0.11, *p* = 0.6, Figure S7A). However, Pleasants *et al.* (2017) point out that these metrics may be biased because few citizen-science records are made from agricultural land, and they provide corrected NABA counts for the period 1999-2014. If we use these corrected counts, there is a much stronger (while not “statistically significant”) correlation between *D. plexippus* occurrence and NABA counts (r = 0.48, *p* = 0.06, Figure S7B).

There was a strong correlation between our *D. plexippus* occurrence and MLMP citizen-science counts of monarch eggs over the period 1999-2014, whether (r = 0.65, *p* < 0.01, Figure S7C) or not (r = 0.66, *p* < 0.01) we used the collection-bias corrections of Pleasants *et al.* (2017).

There was a reasonable (if not significant) correlation between our *D. plexippus* occurrence and estimates of monarch population sizes during the following winter over the period 1994-2014 (r = 0.40, *p* = 0.07, Figure S7D).

Our estimates of *A. syriaca* abundance had a slightly negative correlation with the estimates of *A. syriaca* abundance of Pleasants *et al.* (2017) over the period 1999-2014 (r = -0.35, *p* = 0.2, Figure S7E). However, Pleasants *et al.* (2017) estimated *A. syriaca* abundances from Iowa data alone, while we include *A. syriaca* from across its range in the continental United States. We did not have enough *A. syriaca* records from Iowa during the 1999-2014 period (n = 3) to compare our results more directly.

Overall, previously published data from purpose-built citizen science initiatives predict the size of monarch overwintering populations better than do our data: corrected NABA estimates vs overwintering population size, r = 0.74, *p* < 0.01; MLMP estimates vs overwintering population size, r = 0.55, *p* < 0.05. However, our data are reasonable predictive, supporting their use for the period before 1993 when no other published data sets on monarch or milkweed occurrence are available.

We also tested whether our metrics of *Asclepias* spp. occurrence predicted monarch abundance. Our milkweed occurrence did not predict monarch occurrence over the period 1900-2016 (r = 0.16, *p* = 0.08, Figure S7F). This was still the case when looked at over 5-year bins (r = 0.18, *p* = 0.4, Figure S7G).

Our milkweed occurrence had some mild ability predict the size of the monarch overwintering population the following winter from 1994-2014 (r = 0.34, *p* = 0.14, Figure S7H). However, the milkweed occurrence metric includes many records from states that contribute relatively little to the monarch overwintering population (Flockhart *et al.* 2013). Therefore, we subsequently calculated milkweed occurrence in only those states that contribute the most to the monarch overwintering population. We found that milkweed occurrence in these states does indeed predict the size of the subsequent overwintering population from 1994-2014 (r = 0.45, *p* < 0.05, Figure S7I), albeit not as well as did the purpose-collected data of Pleasants *et al.* (2017) during the period 1999-2014 (r = 0.70, *p* < 0.01).

The weakness of the relationship between occurrence of milkweed and monarchs is perhaps not surprising, as both data sets are relatively noisy at the year-to-year, and even 5-year-to-5-year level, particularly the occurrence of *D. plexippus.* This is likely a combination of sampling error introduced by the method of examining museum collections with natural variation in insect population sizes, as all other metrics of monarch abundance have great amounts of year-to-year variation (Vidal and Rendón-Salinas 2014, Inamine *et al.* 2016). Furthermore, factors beyond milkweed abundance, particularly weather, are known to effect monarch population sizes (Saunders *et al.* 2018). The effect of such other factors on long term trends in monarch population size certainly merits further investigation. However, when views on a decades-to-century time scale, the correspondence between milkweed and monarch occurrence remains striking (Figure 1A).

**Figure S7:**
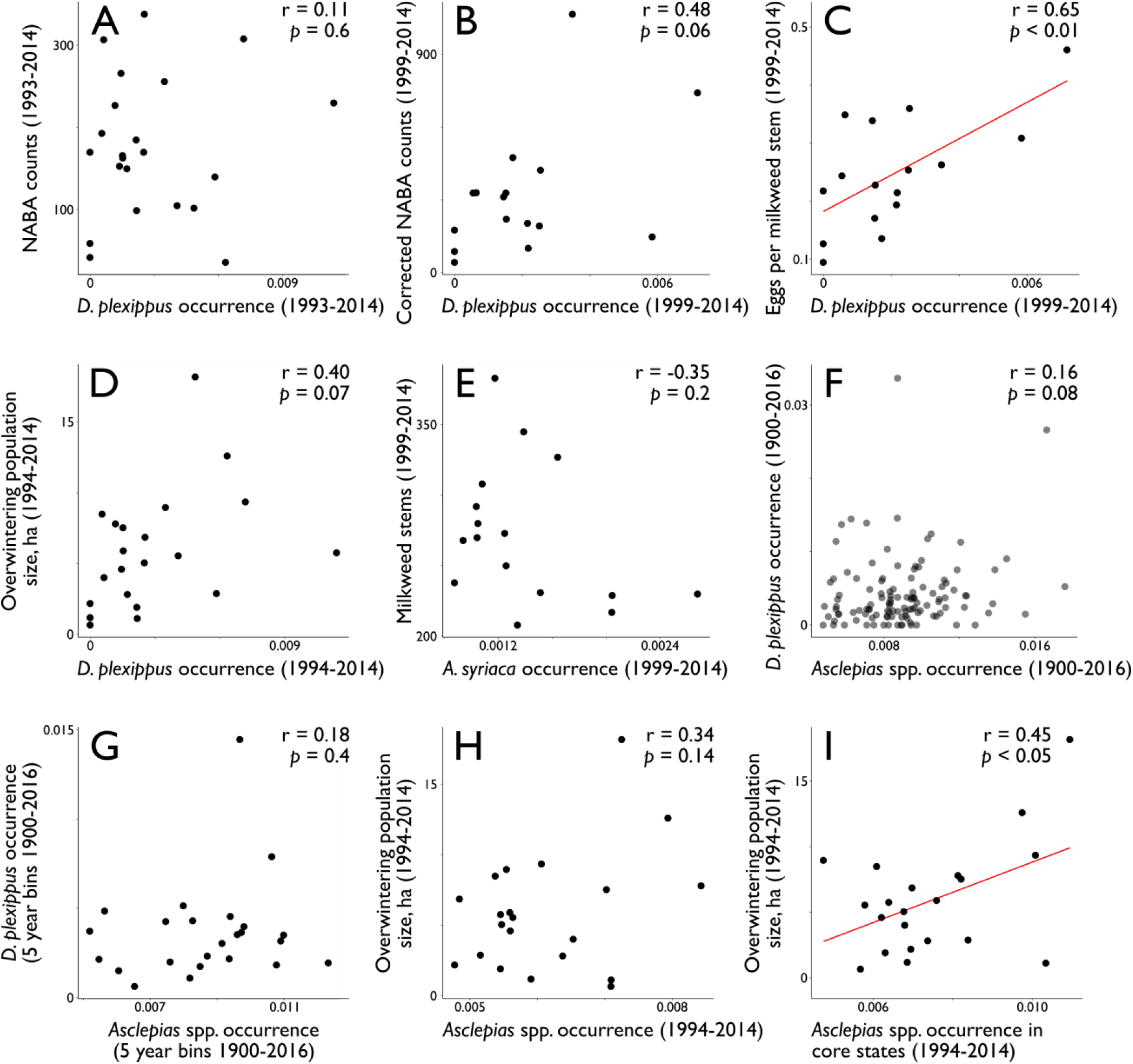
Correlations between various metrics of monarch and milkweed abundance. *Agricultural data*

The nationwide trends for each chosen agricultural variable are shown in Figure S8. We then used state-by-state variation in these characteristics to divide the *A. syriaca* range into six, relatively homogenous regions (Figure S9).

**Figure S8:**
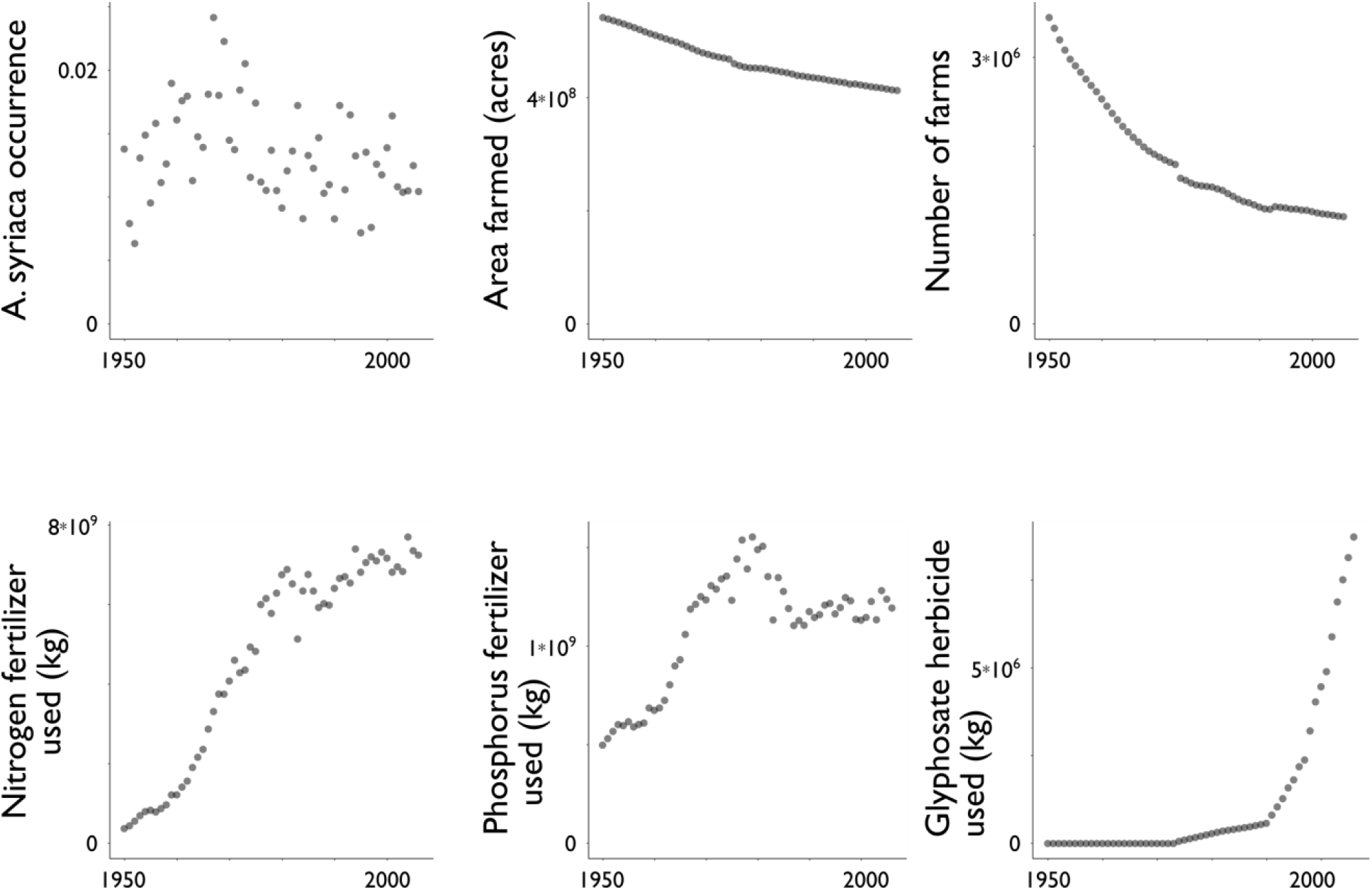
Nationwide trends for chosen agricultural variables. Each point shows the total for all states within the area of our model (see Figure S4) for a single year.

**Figure S9:**
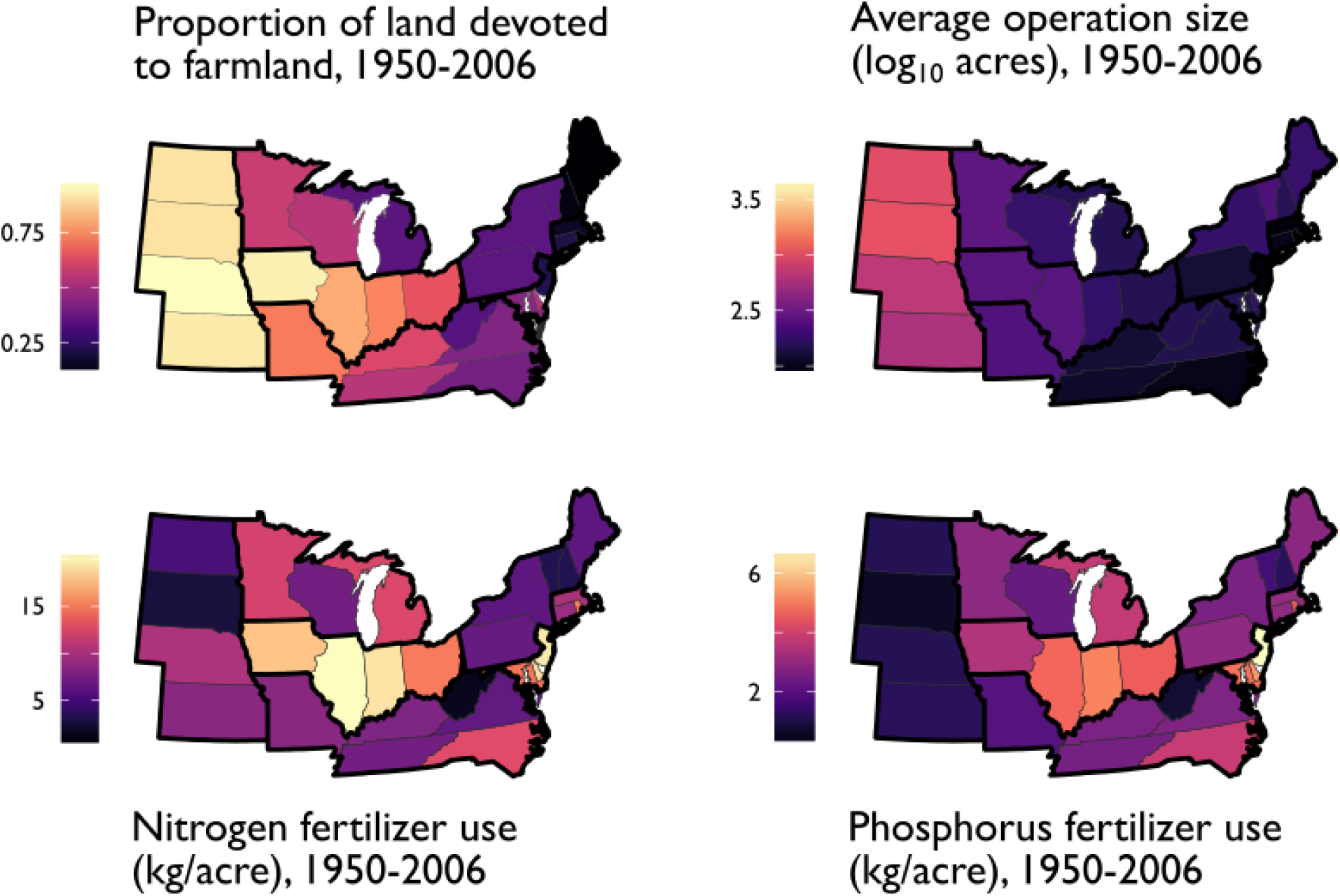
States were grouped into regions (thick black lines) with relatively homogenous agricultural practices.

